# Beyond the fish-*Daphnia* paradigm: testing the potential for *Neoplea striola* (Hemiptera: Pleidae) to cause a trophic cascade in subtropical ponds

**DOI:** 10.1101/2021.04.14.439893

**Authors:** Chase J. Rakowski, Mathew A. Leibold

## Abstract

Trophic cascades, or indirect effects of predators on non-adjacent lower trophic levels, have become paradigmatic in ecology, though they are thought to be stronger in aquatic ecosystems. Most research on freshwater trophic cascades focused on temperate lakes, where fish are present and where *Daphnia* tend to dominate the zooplankton community. These studies identified that *Daphnia* often play a key role in facilitating trophic cascades by linking fish to algae with strong food web interactions. However, *Daphnia* are rare or absent in most tropical and subtropical lowland freshwaters, and fish are absent from small and temporary water bodies, where invertebrates fill the role of top predator. While invertebrate predators are ubiquitous in freshwater systems, most have received little attention in food web research. Therefore, we aimed to test whether trophic cascades are possible in small warmwater ponds where small invertebrates are the top predators and *Daphnia* are absent. We collected naturally occurring plankton communities from small fishless water bodies in central Texas and propagated them in replicate pond mesocosms. We removed zooplankton from some mesocosms, left the plankton community intact in others, and added one of two densities of the predaceous insect *Neoplea striola* to others. Following an incubation period we then compared biomasses of plankton groups to assess food web effects between the trophic levels including whether *Neoplea* caused a trophic cascade by reducing zooplankton. The zooplankton community became dominated by copepods which prefer large phytoplankton and exhibit a fast escape response. Perhaps due to these qualities of the copepods and perhaps due to slow consumption rates by *Neoplea* on key grazers, no evidence for food web effects were found other than somewhat weak evidence for zooplankton reducing large phytoplankton. More research is needed to understand the behavior and ecology of *Neoplea*, but trophic cascades may generally be weak or absent in fishless low-latitude lowland water bodies where *Daphnia* are rare.

## Introduction

An extensive body of literature has demonstrated the importance of direct food web effects, as well as indirect food web effects, or trophic cascades, with much of this work focusing on freshwater ecosystems from an early time. Many of the studies on trophic cascades in freshwater pelagic ecosystems have focused on large-bodied cladocerans, especially *Daphnia*, as the herbivorous prey linking predators to autotrophs (e.g. Carpenter et al. 2001). However, a relative paucity of such studies were performed in tropical and subtropical lowlands, where *Daphnia* is rare (Dumont 1994). Furthermore, the predators that most of these studies focused on were fish, with certain invertebrate taxa such as *Chaoborus* or *Notonecta* receiving some attention while the food web effects of other common and widespread invertebrate predators remain little known (Carpenter et al. 1992, 2001). Understanding the effects of these understudied predaceous invertebrate species is important for understanding their role in ecosystems and for conservation planning, as predators are more threatened with extinction than lower trophic levels (Purvis et al. 2000).

Members of the family Pleidae are small but common heteropteran insects closely related to the family Notonectidae which includes *Notonecta* (backswimmers). Pleids have been much less studied than their larger cousins the notonectids, especially *Notonecta* and *Buenoa*. In this paper we specifically study the pleid *Neoplea striola* (hereafter, *Neoplea*) as our manipulated zooplanktivorous predator. *Neoplea* is a widespread inhabitant of lentic freshwater in Central and North America, and is known to tolerate low oxygen conditions (Gittelman 1975). It is a small insect, with adults measuring 1.5 mm in length. *Neoplea* is an active hunter which uses sight, vibrations, and possibly chemicals to sense its prey (Papacek 2001). They have been shown to attack and consume small zooplankton such as rotifers, mesozooplankton such as *Daphnia*, and even prey as large as dipteran larvae (Gittelman 1974, Hampton and Gilbert 2001, Papacek 2001). However, the effects of *Neoplea* on pond communities are still not fully understood, including whether they can cause trophic cascades.

Here we report the results of a field mesocosm experiment in which we manipulated densities of *Neoplea* and zooplankton to test the effects of both of these trophic groups on plankton composition and biomass, including direct effects on the next trophic level and indirect (trophic cascade) effects of *Neoplea* on phytoplankton. Our plankton communities were composed of freshwater plankton collected locally in central Texas, with no *Daphnia* or other large-bodied cladocerans present. Due to the fast escape response of copepods, we expected *Neoplea* would primarily affect other zooplankton, especially cladocerans and ostracods. We further predicted that zooplankton would reduce the biovolume of total phytoplankton, and that *Neoplea* would cause a trophic cascade, i.e. indirectly increase total phytoplankton biovolume via its suppression of herbivorous zooplankton. However, we found no evidence for food web effects of *Neoplea* and only weak evidence for effects of zooplankton on phytoplankton in the experiment.

## Materials & Methods

### Organism collection

We allowed phytoplankton communities to naturally assemble in two plastic tanks at the University of Texas’ Brackenridge Field Laboratory in Austin, TX for ~3 months. We then mixed 10 L from each tank with 12 L carbonated mineral water to narcotize any zooplankton, filtered this mixture through 45 μm mesh to remove the zooplankton, and mixed well to produce a phytoplankton inoculum. To create a zooplankton inoculum, we collected and mixed zooplankton-rich water from several small fishless water bodies nearby, and concentrated the mixture with a 45 μm mesh. We collected *Neoplea* from small water bodies in Austin, TX.

### Experiment setup and design

We maintained 20 pond communities in 200 L round plastic tanks in an unshaded field at Brackenridge Field Laboratory. We covered the tanks in 1 mm^2^ screens to prevent insect immigration, and maintained constant water depths using float valves. Prior to the experiment, we analyzed total N and P in the water following standard American Public Health Association methods (APHA 1989). We then added NaNO_3_ to bring total N to 14 mg/L N and added NaH_2_PO_4_•H_2_O to bring total P to 1.55 mg/L P. These are the total N and P concentrations in COMBO medium, a eutrophic medium commonly used for culturing plankton (Kilham et al. 1998). Every five or six days thereafter for six weeks, enough of both nutrients were added to each tank to compensate for a 5% daily loss rate from the water column (as per Hall et al. 2004); this same amount of both nutrients was also added immediately following the first sampling (methods described below) after a 22-day pause.

We inoculated each tank with 600 mL of the phytoplankton inoculum. After allowing phytoplankton to reproduce for five days, we added an equal volume of zooplankton inoculum to 15 of the tanks. As there may have been phytoplankton and picoplankton strains in the zooplankton inoculum not represented in the phytoplankton inoculum, we added some of the filtrate left from concentrating the zooplankton inoculum to the other five tanks to ensure all tanks received the same strains. Lastly, after allowing the zooplankton to reproduce for 15 days we added 20 *Neoplea* adults to five tanks with zooplankton and 40 *Neoplea* adults to another five, then added the same amount to the same tanks two days later to bring the totals to 40 and 80. Thus there were four treatments: no zooplankton added (“no zoop.”), zooplankton but no *Neoplea* added (“no *Neoplea*”), zooplankton and 40 *Neoplea* added (“40 *Neoplea*”), and zooplankton and 80 *Neoplea* added (“80 *Neoplea*”). Each treatment was replicated five times for a total of 20 mesocosms arranged randomly.

### Sampling and biomass estimation

We sampled zooplankton and phytoplankton 40 days after adding the *Neoplea*, and then again six days later. To sample zooplankton, we used tube samplers to collect ten whole water column subsamples spread across each tank, and pooled them into a 10 L sample for each tank. We filtered this sample through 65 μm mesh, returning any predators to the tank, and preserved the retained organisms in 10% Lugol’s solution. To sample phytoplankton, we used a 1-cm diameter PVC pipe to collect three whole water column subsamples spread over each tank and pooled them into a 50 mL sample, using a different PVC pipe for each tank. We preserved these phytoplankton samples in 10% Lugol’s solution. We additionally estimated surviving *Neoplea* populations after the second sampling event by using a dipnet to count individuals until we returned three successive empty sweeps.

To estimate biomass of zooplankton taxa, we identified, counted, and measured zooplankton in subsamples such that for each taxon, we counted at least 25 individuals or 10% of the sample – whichever came first – and at least 50 total individuals. We used an ocular micrometer to measure the length of each crustacean and individual *Spirostomum* to the nearest half increment, or 0.24 mm, and to measure the length and width of each rotifer and width of *Spirostomum* to the nearest 0.05 increment, or 0.024 mm. We used length-mass regressions to convert crustacean length to dry mass (McCauley [1984] for *Scapholeberis*, Culver et al. [1985] for copepods, and Anderson et al. [1998] for ostracods). We converted *Spirostomum* dimensions to dry mass by approximating cells as cylinders and assuming a 10:1 biovolume:dry mass ratio. When we were able to identify rotifers to species, we converted the rotifers’ dimensions to dry mass using species-specific equations from the EPA protocol (EPA 2016). For other rotifers, we estimated their biovolume using biovolume equations and then converted to dry mass assuming the same 10:1 biovolume:dry mass ratio (McCauley 1984).

To estimate biovolumes of phytoplankton taxa, we calculated densities of each morphospecies with a hemocytometer. We counted 50 cells or 25 nL of the most common morphospecies – whichever came first – and we counted at least 100 nL for less common taxa. We captured several micrographs of each morphospecies from various tanks and sampling dates, and used ImageJ to measure the cell dimensions of at least 15 cells for all but the rarest morphospecies (Schneider et al. 2012). We then calculated the biovolume of each cell using geometric approximations (Supplemental Table S1).

### Data analysis

Based on previous research (Rakowski et al. 2019), we expected *Neoplea* to have differential effects on zooplankton based on behavior and size. Copepods have faster escape responses than the other zooplankton present, so we analyzed their biomass separately. Then we analyzed the sum of all other zooplankton biomass (“non-copepods”) as we predicted *Neoplea* would primarily affect these other taxa. We also separately analyzed the major groups of non-copepod zooplankton, including cladocerans and ostracods as one group (grouped for similarity in size, morphology, and slow swimming speed), *Spirostomum* as its own distinct category, and lastly rotifers. We expected zooplankton to either affect the whole phytoplankton community, or alternatively to affect only a certain size class, either large or small morphospecies. Therefore we analyzed total phytoplankton biovolume as well as the summed biovolume of larger morphospecies and of smaller morphospecies.

We tested for a difference in *Neoplea* survival between the two densities using a t test. To analyze the effects of zooplankton and *Neoplea* additions on the biomass of plankton groups, we fit a generalized linear mixed model in the gamma family (gamma GLMM) for each plankton grouping using the *lme4* package (Bates et al. 2015). We included tank as a random effect to account for the repeated measures, and fixed effects for zooplankton addition and initial *Neoplea* density. We also fit the nested models excluding *Neoplea* density as well as the null models with neither *Neoplea* density nor zooplankton addition, and used likelihood ratio tests to assess whether including either addition significantly improved model fit. We used one-tailed hypothesis tests for the effect of zooplankton addition on the biomass of zooplankton groups and for the effect of zooplankton and *Neoplea* addition on total phytoplankton. In all other cases we used two-tailed hypothesis tests since it was conceivable that a reduction of one plankton group could have benefited another plankton group in the same trophic level. For example, zooplankton could conceivably have had a negative effect on large phytoplankton and a positive effect on small phytoplankton. All analyses were performed in R v. 3.5.3 (R Core Team 2017).

## Results

On average *Neoplea* survived at a rate of 74%, with no significant difference in survival between the two densities (d.f. = 7.98, *t* = −0.518, *p* = 0.618). No evidence was found of *Neoplea* reproduction during the experiment.

By the sampling dates, the zooplankton community had become dominated by copepods (Supplemental Table S2). However, cladocerans, *Spirostomum*, and rotifers all individually composed at least 15% of the zooplankton mass on average in one or more treatments (Fig. 1). A small amount of zooplankton, mostly copepods, became established in the control tanks receiving no zooplankton (Fig. 1, Fig. 2). The zooplankton additions were successful in sustaining a significant increase in biomass of all zooplankton groups (Table 1, Fig. 2). However, *Neoplea* additions had no significant effects on the biomass of any zooplankton group (Fig. 2, Table 1). Mean copepod mass was higher and the mean proportion of rotifers was lower the more *Neoplea* were added (Fig. 1), but neither copepod nor rotifer mass was significantly affected by *Neoplea* (Fig. 2, Table 1).

**Figure 1:**
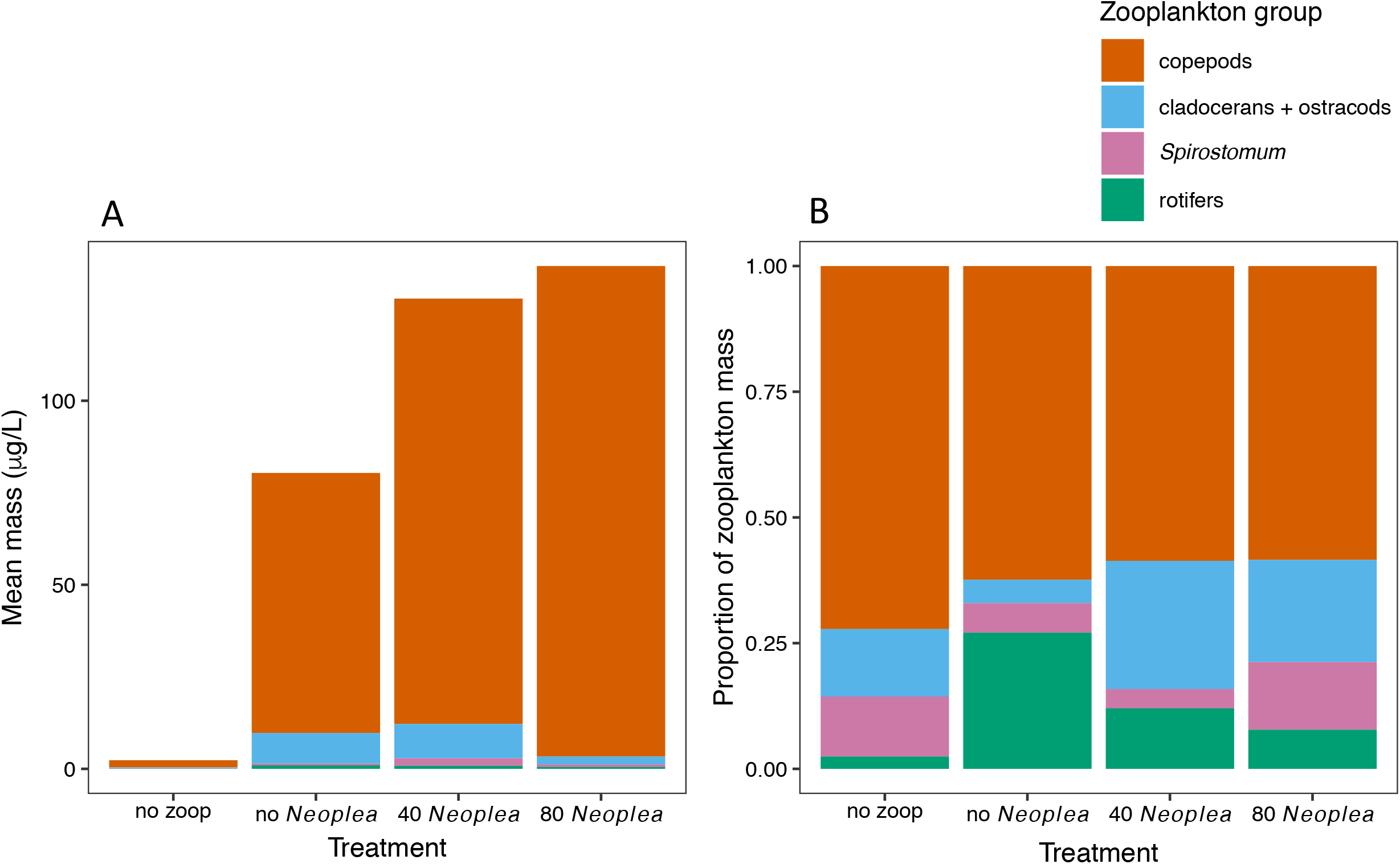
Zooplankton composition across treatments. (A) Stacked bar chart representing mean dry mass of zooplankton groups by treatment. (B) Stacked bar chart representing mean proportions of total zooplankton dry mass each group comprises, by treatment.

**Figure 2:**
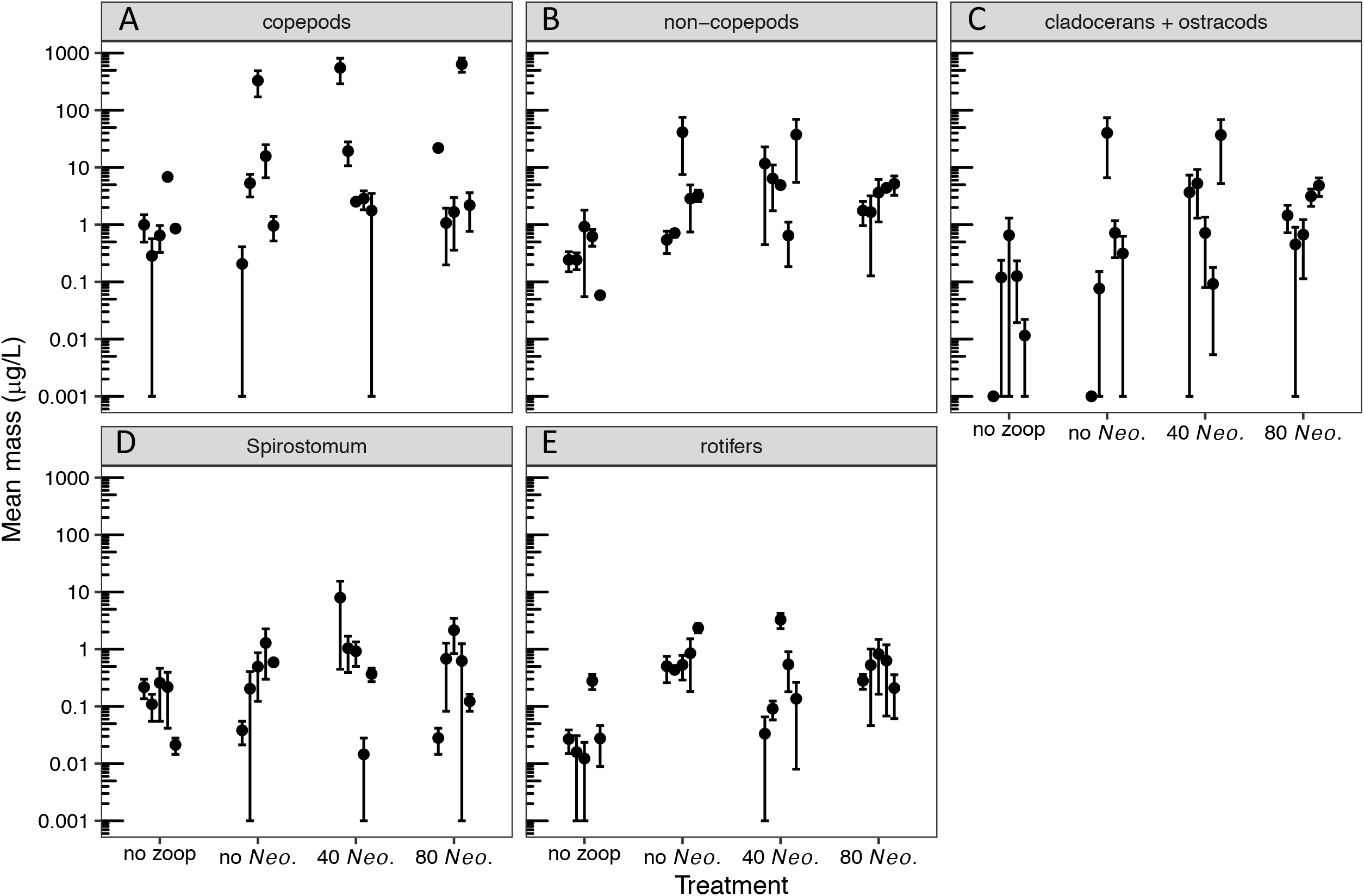
Dry mass of zooplankton groups by treatment. Dots represent means of individual tanks, and error bars represent temporal standard errors. (A) copepods, (B) non-copepod zooplankton, (C) cladocerans and ostracods, (D) *Spirostomum*, and (E) rotifers. Note that 0.001 was added to all values to allow plotting on a log scale.

**Table 1:**
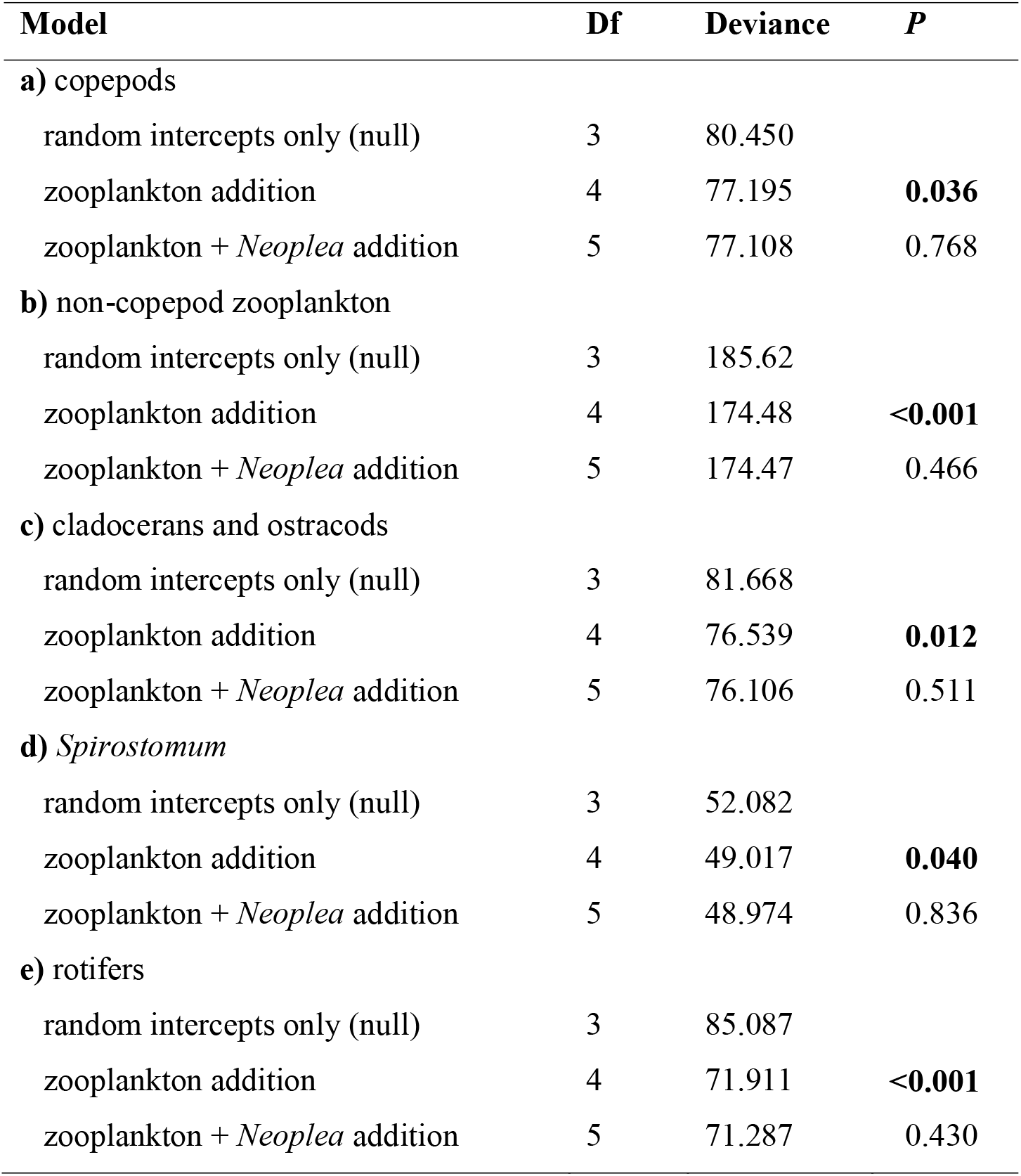
Results of likelihood ratio tests comparing nested GLMMs for biomass of zooplankton groups. Displayed are the degrees of freedom (Df), deviance (inverse goodness of fit), and *P* value for comparison against the model above, for biomass of **a)** copepods, **b)** non-copepod zooplankton, **c)** cladocerans and ostracods, **d)** *Spirostomum*, and **e)** rotifers. *P* values <0.05 are bolded.

**Table 2:** Coefficient estimates for GLMMs analyzing the effect of zooplankton addition on the biomass of zooplankton groups. Estimates are displayed along with their standard errors (SE) and natural exponential functions [“Exp(estimate)”] for **a)** copepods, **b)** non-copepod zooplankton, **c)** cladocerans and ostracods, **d)** *Spirostomum*, and **e)** rotifers. The natural exponential functions of estimates can be interpreted as multiplicative effects (e.g. zooplankton addition resulted in a 7.66-fold increase in copepod mass). The effect of *Neoplea* addition is not included due to likelihood ratio tests indicating the term was not significant for any zooplankton group.

| Parameter | Estimate | SE | Exp(estimate) |
| --- | --- | --- | --- |
| **a) copepods** |  |  |  |
| intercept | 0.382 | 0.932 |  |
| zooplankton addition | 2.036 | 1.077 | 7.660 |
| **b) non-copepod zooplankton** |  |  |  |
| intercept | -1.184 | 0.559 |  |
| zooplankton addition | 2.519 | 0.646 | 12.416 |
| **c) cladocerans and ostracods** |  |  |  |
| intercept | -1.346 | 0.704 |  |
| zooplankton addition | 2.003 | 0.816 | 7.411 |
| **d) *Spirostomum*** |  |  |  |
| intercept | -1.257 | 0.459 |  |
| zooplankton addition | 1.018 | 0.538 | 2.768 |
| **e) rotifers** |  |  |  |
| intercept | -3.159 | 0.510 |  |
| zooplankton addition | 2.572 | 0.588 | 13.092 |

The phytoplankton community became dominated by ovoid single-celled green algae, with pennate diatoms contributing the next most biovolume in tanks with no zooplankton added and *Oocystis* contributing the next most biovolume in tanks with 80 *Neoplea* added (Fig. 3, Supplemental Table S1). While the average total phytoplankton biovolume with zooplankton was less than half of the average without zooplankton, there was no significant effect of zooplankton addition on total phytoplankton biovolume due to large variation within treatments (Fig. 3, Fig. 4A, Table 3A, Table 4A). When the largest morphospecies were taken together, including the larger ovoid chlorophytes, pennate diatoms, and *Oocystis*, zooplankton reduced their summed biovolume marginally significantly by 77.9% (Fig. 4B, Table 3B, Table 4B). On the other hand, summed biovolume of the smaller morphospecies (small ovoid chlorophytes, *Chlorella*, *Selenastrum*, and photosynthetic picoplankton) was not significantly impacted by zooplankton addition (Fig. 4C, Table 3C, Table 4C). Similarly, *Neoplea* addition had no significant effects on the biovolume of any phytoplankton grouping (Fig. 4, Table 3).

**Figure 3:**
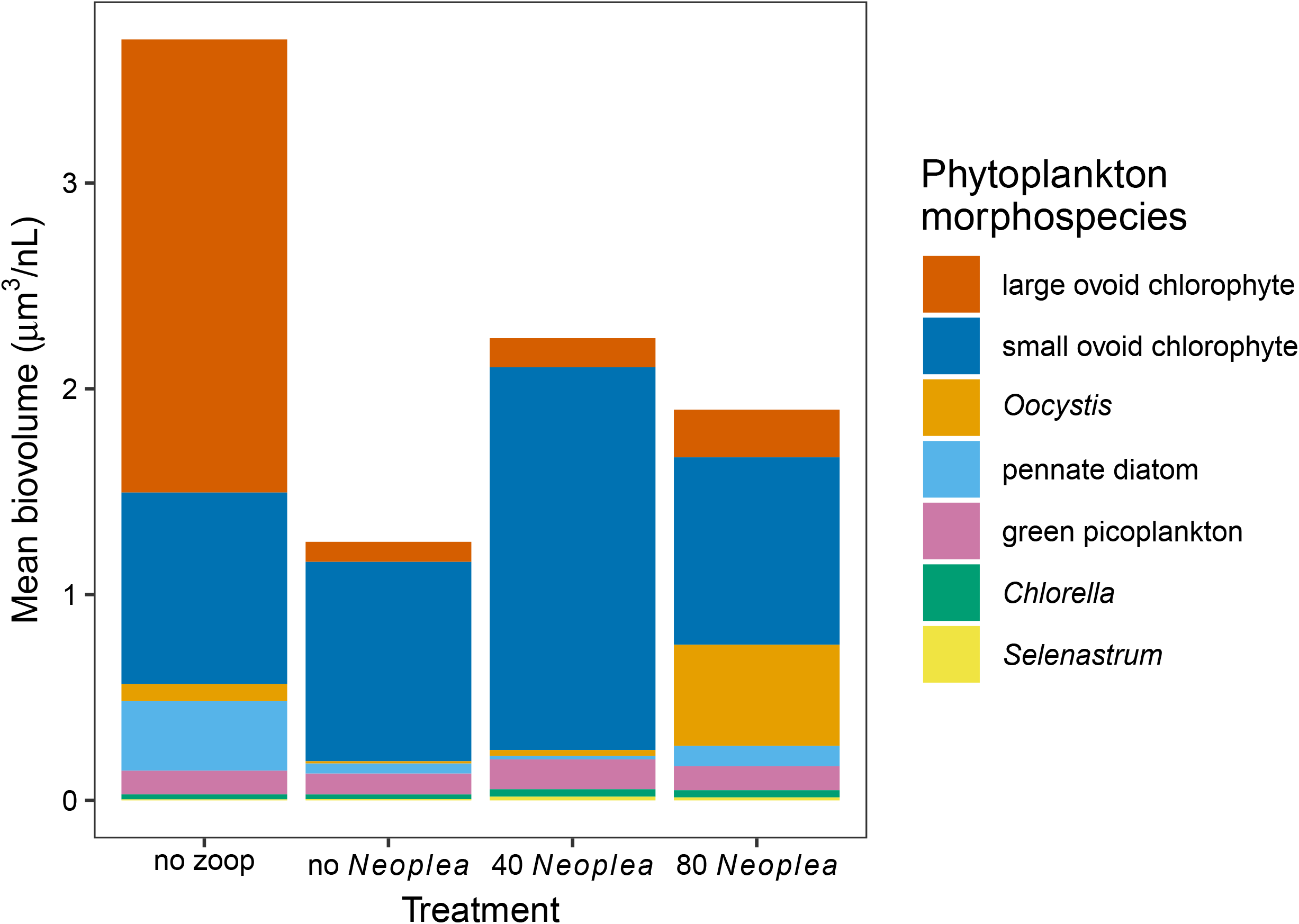
Phytoplankton composition across treatments. Stacked bar chart of mean biovolume of phytoplankton morphospecies, by treatment.

**Figure 4:**
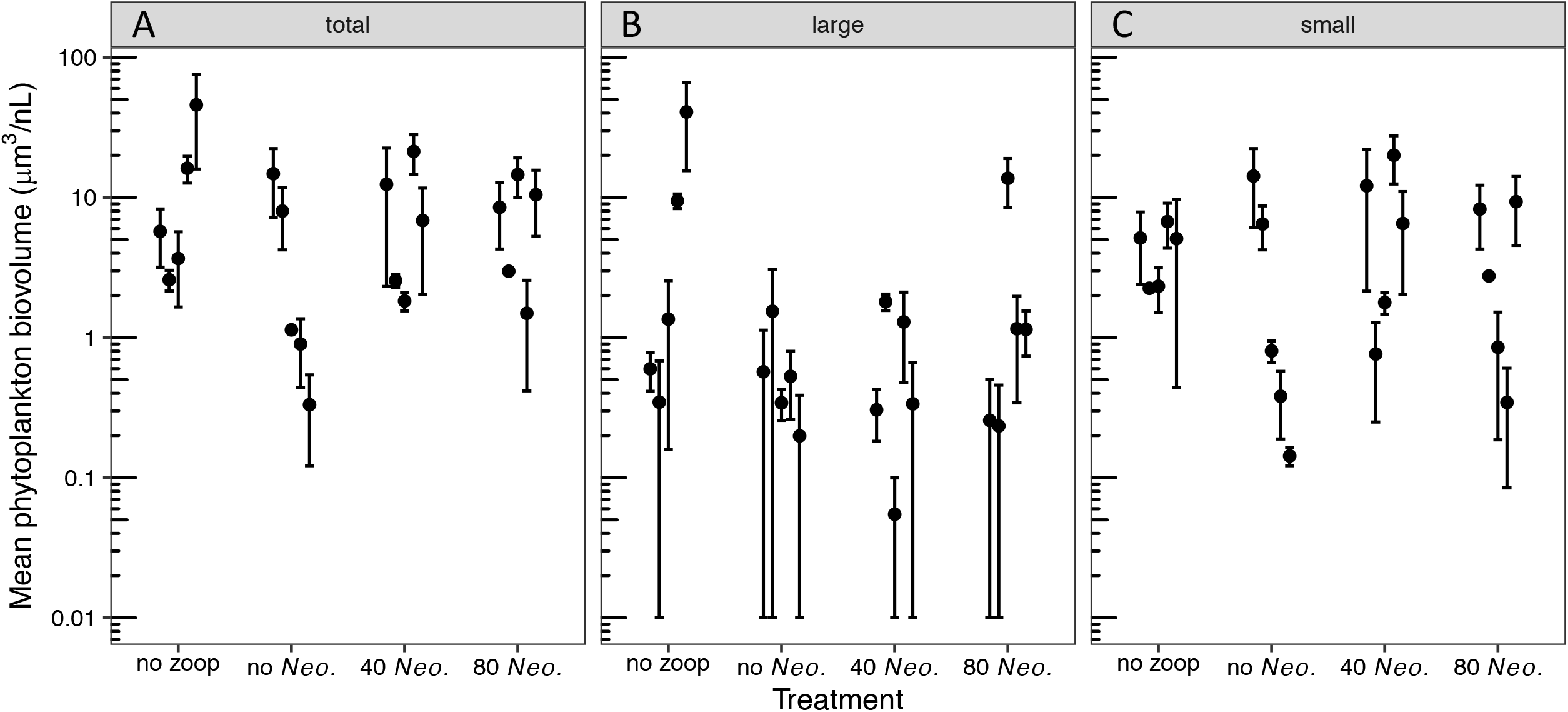
Phytoplankton biovolume by treatment. Dots represent means of individual tanks, and error bars represent temporal standard errors. (A) total phytoplankton; (B) large phytoplankton (larger ovoid chlorophytes, *Oocystis*, and pennate diatoms); and (C) small phytoplankton (smaller ovoid chlorophytes, picoplankton, *Chlorella*, and *Selenastrum*). Note that 0.01 was added to all values to allow plotting on a log scale.

**Table 3:** Results of likelihood ratio tests comparing nested GLMMs for biovolume of phytoplankton groupings. Displayed are the degrees of freedom (Df), deviance (inverse goodness of fit), and *P* value for comparison against the model above, for biovolume of **a)** total phytoplankton, **b)** larger phytoplankton (large ovoid chlorophytes, *Oocystis*, and pennate diatoms), and **c)** smaller phytoplankton (small ovoid chlorophytes, green picoplankton, *Chlorella*, and *Selenastrum*).

| Model | Df | Deviance | *P* |
| --- | --- | --- | --- |
| **a) total phytoplankton** |  |  |  |
| random intercepts only (null) | 3 | 786.53 |  |
| zooplankton addition | 4 | 785.50 | 0.155 |
| zooplankton + *Neoplea* addition | 5 | 783.90 | 0.103 |
| **b) larger phytoplankton** |  |  |  |
| random intercepts only (null) | 3 | 101.79 |  |
| zooplankton addition | 4 | 98.06 | 0.053 |
| zooplankton + *Neoplea* addition | 5 | 97.32 | 0.389 |
| **c) smaller phytoplankton** |  |  |  |
| random intercepts only (null) | 3 | 749.71 |  |
| zooplankton addition | 4 | 749.25 | 0.500 |
| zooplankton + *Neoplea* addition | 5 | 748.87 | 0.535 |

**Table 4:**
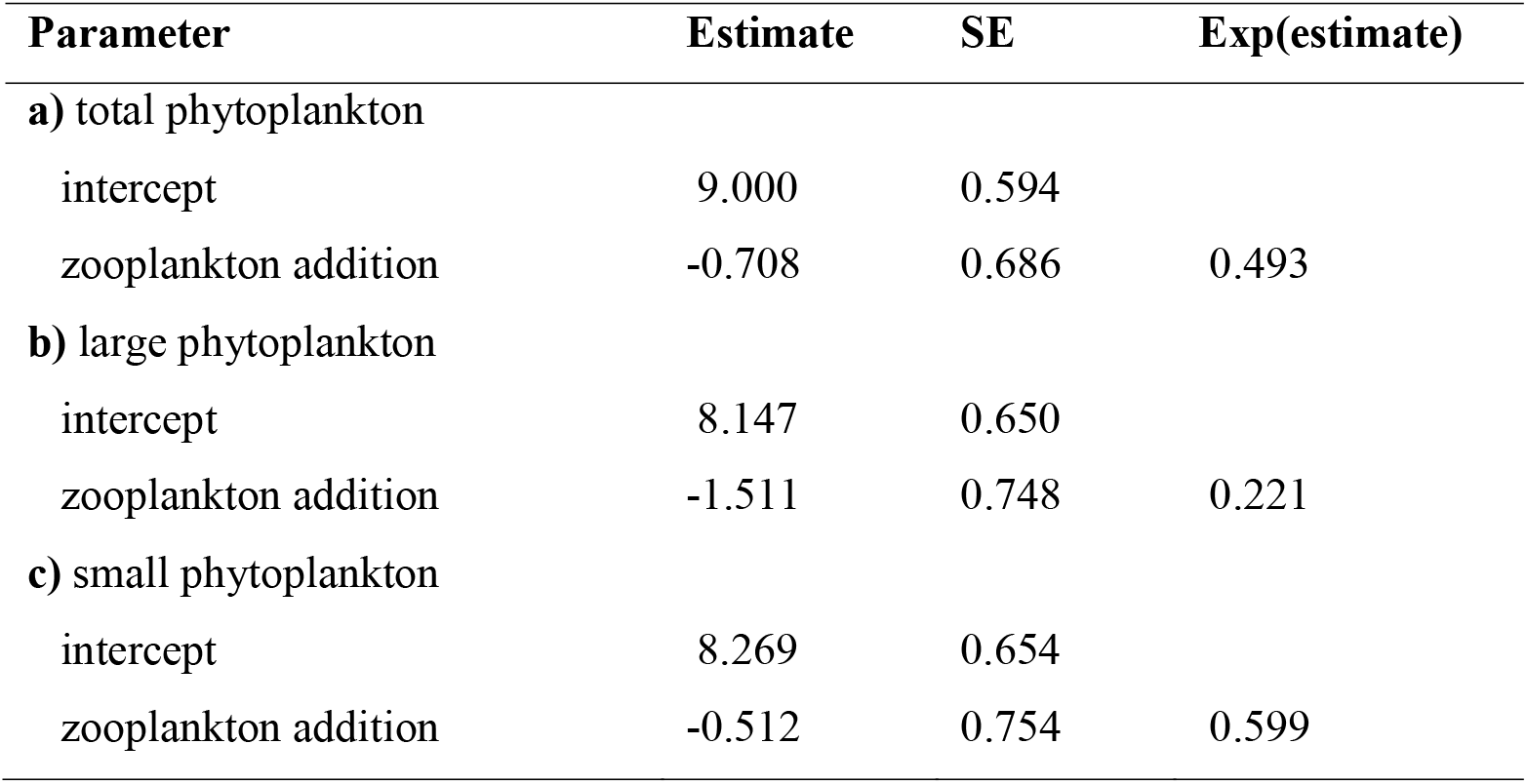
Coefficient estimates for GLMMs analyzing the effect of zooplankton addition on the biovolume of phytoplankton groupings. Estimates are displayed along with their standard errors (SE) and natural exponential functions [“Exp(estimate)”] for **a)** total phytoplankton, **b)** larger phytoplankton (large ovoid chlorophytes, *Oocystis*, and pennate diatoms), and **c)** smaller phytoplankton (small ovoid chlorophytes, green picoplankton, *Chlorella*, and *Selenastrum*). The natural exponential functions of estimates can be interpreted as multiplicative effects (e.g. zooplankton addition reduced large phytoplankton to 22.1% of the control biovolume). The effect of *Neoplea* addition is not included due to likelihood ratio tests indicating the term was not significant for any phytoplankton grouping.

## Discussion

Our results show that *Neoplea* had no demonstrable effect on the plankton in this experiment. Furthermore, the copepod-dominated zooplankton only caused a marginal reduction of larger phytoplankton taxa. Any effect of *Neoplea* on phytoplankton composition or biomass would have likely been mediated by an effect on zooplankton composition or biomass, as *Neoplea* are carnivorous. Therefore it is unsurprising there was no indirect effect of *Neoplea* on phytoplankton, or trophic cascade, considering there were only weak or nonexistent direct effects between the three trophic levels.

There are several potential reasons why *Neoplea* did not reduce the biomass of any zooplankton groups in our experiment. First, it is possible *Neoplea*’s predation rate was similar enough to the reproduction rate of its prey that prey populations did not significantly change over the course of the 46-day experiment, a scenario most likely if *Neoplea* preferred prey with short generation times such as cladocerans. While a few studies observed predation by *Neoplea* on various prey taxa, we are aware of only one study that measured the effect of *Neoplea* on the biomass of prey taxa over time (Rakowski et al. 2019). In that study *Neoplea* had strong effects on non-copepod zooplankton, which were dominated by fast-reproducing cladocerans, on the short term (under one cladoceran generation). In contrast, *Neoplea* had weak effects on a longer term (~six cladoceran generations) in the absence of other predators. Interestingly however, when the larger predator *Notonecta* was also present, *Neoplea* acted synergistically with *Notonecta* to suppress the non-copepod zooplankton over this longer time frame. Therefore it may be that *Neoplea* exhibits a slow predation rate and prefers prey with short generation times, with the effect that over the long term it can have a cryptic food web effect not apparent unless other predators are present.

Other potential reasons for the lack of an effect of *Neoplea* on zooplankton relate to other aspects of the predaceous insect’s behavior and to the composition of potential prey. It is possible that *Neoplea* consumed chironomid larvae, which were common in the mesocosms but not effectively sampled due to their benthic nature and our sampling methods which targeted plankton. *Neoplea* is known to attack and consume chironomid larvae (Papacek 2001). Indeed, the *Neoplea* were most commonly observed clinging to the sides and bottom of the mesocosms, putting them in close proximity to the chironomids. In nature, *Neoplea* is normally found clinging to submerged vegetation (Gittelman 1974). However, it was impossible to observe the *Neoplea* throughout most of the experiment due to the high density of phytoplankton. This high turbidity also may have made it difficult for the *Neoplea* to hunt effectively, as they can detect prey visually and are most commonly found in clear waters, though they can also detect prey by tactile and possibly chemosensory methods (Gittelman 1974, Papacek 2001). When the *Neoplea* did swim through the water column, they would have mostly encountered copepods, the largest and dominant group of zooplankton. However, even in clear water copepods are relatively resistant to predation due to their fast escape response. Among copepods, diaptomids have an especially fast escape response, and the diaptomid *Arctodiaptomus* dominated our mesocosms. Therefore it may not be surprising that the *Neoplea* were unsuccessful in suppressing the zooplankton in the experiment. While other more easily captured prey were also present, it may have been difficult for the *Neoplea* to encounter these rare prey among all the copepods in a turbid environment.

The dominance of copepods in the zooplankton community may also explain the weak effects of zooplankton on phytoplankton in the experiment. Unlike *Daphnia*, copepods do not generally impose strong top-down control on community phytoplankton biomass, due in part to their selective grazing on larger phytoplankton (Sommer and Sommer 2006). Indeed, the copepod-dominated zooplankton only reduced larger phytoplankton, and even this effect was marginal, fitting the general understanding of copepods’ top-down effects on phytoplankton in ecosystems with copepod-dominated herbivore communities such as open oceans (Sommer and Sommer 2006). Such weak top-down control of crustacean zooplankton on phytoplankton may be more pervasive at low latitudes, as *Daphnia* is largely absent from tropical and subtropical lowland freshwaters and copepods are instead more likely to dominate (Dumont 1994, Havens and Beaver 2011). In combination with the ability of copepods to evade predation much more easily than *Daphnia*, it appears likely that trophic cascades mediated by zooplankton are less common in warm, low-latitude lakes and ponds than in their colder counterparts (Rejas et al. 2005).

The food web ecology of *Neoplea* and pleids generally will remain unclear without further research. More behavioral research on these diminutive predators is needed to better understand their hunting habits, such as where in the habitat they make most of their captures, their relative dependence on different senses for prey detection, and their relative preference for various prey. Experimental work is needed to better quantify the ecology of pleids, and these experiments will likely benefit from better replicating their preferred habitat of clear, still water with submerged vegetation. A higher population density may be necessary for their food web effects to be apparent when they are the sole predator.

## Conclusions

We predicted that *Neoplea* would suppress non-copepod zooplankton, and that zooplankton would suppress phytoplankton, resulting in a trophic cascade where *Neoplea* indirectly increased phytoplankton biomass. Instead, *Neoplea* had no significant effect on plankton biomass or composition in this field mesocosm experiment. The zooplankton, dominated by copepods and lacking *Daphnia* as is typical of lowland tropical and subtropical lakes and ponds, only weakly reduced larger phytoplankton. While our data cannot definitively explain these weak effects, they could have resulted from *Neoplea* consuming prey able to reproduce quickly enough to make up for losses to predation, from *Neoplea* consuming benthic prey which was not effectively sampled, or from the dominance of copepods which are adept at evading capture and which selectively feed on larger phytoplankton. This study suggests that lentic ecosystems dominated by *Neoplea* and copepods may be characterized by weak top-down control. This represents another example of a weak or non-existent trophic cascade in lowland tropical or subtropical lentic freshwater, which appears to be much more common in these systems than in colder lakes and ponds. However, more research will be needed to achieve a clearer understanding of the ecological impacts of pleids. A better understanding of the ecology of these and other understudied invertebrate predators will be important for conservation planning, as predators are more threatened with extinction than lower trophic levels, a pattern that does not only apply to charismatic megafauna.

## Acknowledgements

Thanks to L. A. Sekula for help processing samples, to S. Manning for providing laboratory space, to R. Deans for help with insect collection and experiment setup, to S. Duchicela for help with experimental setup, to D. Correa for help with nutrient analysis, and to D. Nobles for providing equipment. This research was supported by the Department of Integrative Biology at the University of Texas at Austin and was made possible by the facilities at Brackenridge Field Laboratory.

## Notes

### Competing Interest Statement

The authors have declared no competing interest.

### Summary of Updates

Figure legends and tables added to manuscript. Supplemental tables added. Minor revisions made to manuscript text.

## References

Anderson, D. H., S. Darring, and A. C. Benke. 1998. Growth of crustacean meiofauna in a forested floodplain swamp: implications for biomass turnover. Journal of the North American Benthological Society 17:21–36.

APHA. 1989. Standard methods for the examination of water and wastewater, 17th edn. American Public Health Association, Washington, D.C.

Bates, D., M. Maechler, B. Bolker, and S. Walker. 2015. Fitting linear mixed-effects models using lme4. Journal of Statistical Software 67:1–48.

Carpenter, S., J. Cole, J. Hodgson, J. Kitchell, M. Pace, D. Bade, K. Cottingham, T. Essington, J. Houser, and D. Schindler. 2001. Trophic cascades, nutrients, and lake productivity: whole-lake experiments. Wiley, Hoboken, NJ.

Carpenter, S., C. Kraft, R. Wright, H. Xi, P. Soranno, and J. Hodgson. 1992. Resilience and resistance of a lake phosphorus cycle before and after food web manipulation. Univ Chicago Press, Chicago, IL.

Culver, D. A., M. M. Boucherle, D. J. Bean, and J. W. Fletcher. 1985. Biomass of freshwater crustacean zooplankton from length-weight regressions. Canadian Journal of Fisheries and Aquatic Sciences 42:1380–1390.

Dumont, H. 1994. On the diversity of Cladocera in the tropics. Springer, Dordrecht, Netherlands.

EPA. 2016. Standard operating procedure for zooplankton analysis (LG403, Revision 07). Environmental Protection Agency, Washington, D.C.

Gittelman, S. H. 1974. The habitat preference and immature stages of Neoplea striola (Hemiptera: Pleidae). Journal of the Kansas Entomological Society 47:491–503.

Gittelman, S. H. 1975. Physical gill efficiency and winter dormancy in the pigmy backswimmer, Neoplea striola (Hemiptera: Pleidae). Annals of the Entomological Society of America 68:1011–1017.

Hall, S. R., M. A. Leibold, D. A. Lytle, and V. H. Smith. 2004. Stoichiometry and planktonic grazer composition over gradients of light, nutrients, and predation risk. Ecology 85:2291–2301.

Hampton, S. E., and J. J. Gilbert. 2001. Observations of insect predation on rotifers. Pages 115–121 *in* L. Sanoamuang, H. Segers, R. J. Shiel, and R. D. Gulati, editors. Rotifera IX. Springer Netherlands, Dordrecht.

Havens, K. E., and J. R. Beaver. 2011. Composition, size, and biomass of zooplankton in large productive Florida lakes. Hydrobiologia 668:49–60.

Kilham, S. S., D. A. Kreeger, S. G. Lynn, C. E. Goulden, and L. Herrera. 1998. COMBO: a defined freshwater culture medium for algae and zooplankton. Hydrobiologia 377:147–159.

McCauley, E. 1984. The estimation of the abundance and biomass of zooplankton in samples. Pages 228–265 *in* J. A. Downing and F. H. Rigler, editors. A Manual for the Assessment of Secondary Productivity in Fresh Waters. Blackwell Scientific Publishers.

Papacek, M. 2001. Small aquatic and ripicolous bugs (Heteroptera: Nepomorpha) as predators and prey: the question of economic importance. European Journal of Entomology 98:1–12.

Purvis, A., J. L. Gittleman, G. Cowlishaw, and G. M. Mace. 2000. Predicting extinction risk in declining species. Proceedings of the Royal Society of London. Series B: Biological Sciences 267:1947–1952.

Rakowski, C. J., C. E. Farrior, S. R. Manning, and M. A. Leibold. 2019. Predator complementarity dampens variability of phytoplankton biomass in a diversity-stability trophic cascade. bioRxiv, <https://doi.org/10.1101/851642>.

R Core Team. 2017. R: a language and environment for statistical computing. R Foundation for Statistical Computing, Vienna, Austria.

Rejas, D., S. Declerck, J. Auwerkerken, P. Tak, and L. De Meester. 2005. Plankton dynamics in a tropical floodplain lake: fish, nutrients, and the relative importance of bottom-up and top-down control. Wiley, Hoboken, NJ.

Schneider, C. A., W. S. Rasband, and K. W. Eliceiri. 2012. NIH Image to ImageJ: 25 years of image analysis. Nature Methods 9:671–675.

Sommer, U., and F. Sommer. 2006. Cladocerans versus copepods: the cause of contrasting top–down controls on freshwater and marine phytoplankton. Oecologia 147:183–194.

