## Supplemental Table 1 for "Beyond the fish-*Daphnia* paradigm: testing the potential for *Neoplea striola* (Hemiptera: Pleidae) to cause a trophic cascade in subtropical ponds"

**Table S1:**

**Phytoplankton morphospecies/taxa and their cell shape and mean cell biovolume.**

Taxa are listed in descending order of dominance defined by mean biovolume across all samples.

| **Morphospecies/taxon** | **Shape** | **Mean cell biovolume (μm^3^)** | **Group** |
| --- | --- | --- | --- |
| small ovoid chlorophyte | spheroid | 10.5 | small |
| large ovoid chlorophyte | spheroid | 70.9 | large |
| *Oocystis* | spheroid | 75.1 | large |
| pennate diatom | pennate (rectangular prism tapering to points at each end) | 56.7 | large |
| green picoplankton | sphere | 2.11 | small |
| green round (*Chlorella*) | sphere | 4.95 | small |
| *Selenastrum* | curved cylinder ending in 2 half-spheres | 11.1 | small |
