## Supplemental Table 2 for "Beyond the fish-*Daphnia* paradigm: testing the potential for *Neoplea striola* (Hemiptera: Pleidae) to cause a trophic cascade in subtropical ponds"

**Table S2:**

**Zooplankton taxa, groupings, and dry biomass across all samples.**

Taxa (and stages) are listed in descending order of dominance, determined by median biomass or by mean biomass when the median is 0. Eight taxa not listed here, all rotifers, were rare and represented a negligible contribution to total zooplankton mass.

| **Taxon/stage** | **Classification** | **Grouping** | **Median mass (μg/L)** | **Mean mass (μg/L)** |
| --- | --- | --- | --- | --- |
| *Arctodiaptomus dorsalis* (copepodites & adults) | Copepoda | copepods | 1.13 | 71.5 |
| *Microcyclops varicans* (copepodites & adults) | Copepoda | copepods | 0.412 | 1.61 |
| copepod nauplii | Copepoda | copepods | 0.332 | 7.23 |
| *Spirostomum* sp. 1 - large girth | Ciliophora | *Spirostomum* | 0.230 | 0.839 |
| *Scapholeberis kingi* | Cladocera | cladocerans + ostracods | 0.165 | 4.85 |
| small ploimid | Rotifera | rotifers | 0.0446 | 0.298 |
| *Monostyla quadridentata* | Rotifera | rotifers | 0.00701 | 0.196 |
| ostracod | Ostracoda | cladocerans + ostracods | 0 | 0.0859 |
| cladoceran embryo | Cladocera | cladocerans + ostracods | 0 | 0.0414 |
| *Lecane luna* | Rotifera | rotifers | 0 | 0.0409 |
| *Spirostomum* sp. 2 - small girth | Ciliophora | *Spirostomum* | 0 | 0.0284 |
| *Monostyla lunaris* | Rotifera | rotifers | 0 | 0.0189 |
| bdelloid | Rotifera | rotifers | 0 | 0.00872 |
| *Monostyla copeis* | Rotifera | rotifers | 0 | 0.00572 |
| *Monostyla closterocerca* | Rotifera | rotifers | 0 | 0.00409 |
